# Development of an In Vitro Test for the Optimization of Drug Diffusion in Pediatric Solid Tumors

**DOI:** 10.1101/2022.05.23.493070

**Authors:** Rachel Ivy, Alissa Hendricks-Wenger, Lyndon Kennedy, Anna Jones, Deanna Riley, Ashley Handy, Elizabeth D. Barker

**Affiliations:** Department of Mechanical, Aerospace, and Biomedical Engineering, The University of Tennessee at Knoxville, Knoxville, Tennessee; Lincoln Memorial University - DeBusk College of Osteopathic Medicine, Knoxville, Tennessee

## Abstract

There is significant long-term morbidity and mortality associated with the treatment of childhood cancer, and the risk of these effects continues to increase years after completion of therapy. Among childhood cancer survivors the cumulative incidence of a chronic health condition is 99% within 50 years of the original cancer diagnosis. There is a high risk for severe, disabling, or life-threatening chronic condition caused by the chemotherapy used to treat the initial malignancy. Current standards for determining chemotherapy dosage to treat solid tumor malignancies of pediatric patients is based on several factors, including the patient’s surface area, age, weight, and height. To reduce the long-term effects of chemotherapy in pediatric patients our group is focused on developing novel local drug delivery systems to treat solid tumors and minimize systemic effects. The aim of the current study is to develop an *in vitro* method to quantify drug diffusion through tumor tissue that will allow us to optimize the dose required to treat solid tumor malignancies *in vivo*. We hope by modeling the significant parameters that influence drug penetration of chemotherapy drugs, we can facilitate the development of innovative drug delivery methods and more effective administration of anticancer agents to better treat pediatric malignancies and improve both short-term and long-term outcomes for childhood cancer.

## Background

Among childhood cancer survivors, 99% develop chronic conditions from the toxicity of the chemotherapy used to treat their initial malignancy [1-4]. These chronic conditions can be severe, disabling, or even life-threatening and can manifest years after completion of therapy. Brain and CNS tumors are responsible for 20% of all childhood cancer deaths and survivors of these malignancies are at highest risk for long-term and late effects from their treatment. Current standards for determining chemotherapy dosage to treat solid tumor malignancies of pediatric patients is based on several factors, including the patient’s surface area, age, and weight [5]. However, due to the physiological limitations of systemically delivered drugs to CNS tumors there is a need to develop novel delivery methods [3]. To improve treatment efficacy and prevent long-term and late-effects in pediatric cancer patients, there are many groups working to develop novel local drug delivery systems to treat solid tumors [6]. We expect the dose for these types of drug delivery systems to be determined by the characteristics of the tumor, such as volume and density, and not the size of the patient.

One chemotherapy utilized in the treatment of pediatric CNS tumors is the tumoricidal drug doxorubicin (DOX). A chemosensitivity analysis of pediatric glioblastoma, anaplastic astrocytoma, and other high-grade gliomas showed doxorubicin-induced significant cell death [7]. However, even with potent *in vitro* cytotoxicity and the application of various drug delivery methods, DOX is not able to achieve significant tumor reduction *in vivo* [8-14]. To address this need, our group and others are working to develop local drug delivery methods to overcome systemic side effects and maximize local drug concentration [15-17].

The aim of the current project is to develop an *in vitro* testing system to optimize dose from local drug delivery systems to treat solid tumor malignancies. Through this pilot project, our current goal is to establish an *in vitro* method for mapping drug penetration of chemotherapy agents, using DOX as a representative, released from the drug delivery system over time. By simulating the penetration distance of chemotherapies *in vitro*, we can facilitate the development of innovative drug delivery methods and more effective administration of anticancer agents to better treat pediatric malignancies and improve both short-term and long-term outcomes for childhood cancer.

## Methods

### Synthesis of Agarose Brain Tissue Phantoms

A 0.6% agarose gel model for healthy brain tissue was utilized to evaluate drug release and distribution, because of its comparable mechanical properties to healthy brain tissue [18]. We combined DMSO, agarose powder, and TBE Buffer to create 0.6% agarose gel with the method adopted by Chen, Z.-J., et al. It was then heated using the Anton Parr synthetic microwave to 90°C for 5 minutes to reduce concentration of dissolved gasses. Then it was cooled to 50°C and pipetted into a 6 well plate. A total of 8mL of the solution was pipetted into each plate and it was then placed on the lab bench to allow time for the gel to form.

### Fluorescent Quantification of Distribution

DOX, an auto fluorescent drug, was used as a model compound to stablish this *in vitro* diffusion evaluation method. Monitoring of the DOX diffusion was performed using a PerkinElmer IVIS Lumina K to measure the autofluorescence produced by the drug.

To quantify the diffusion of drug over time, fluorescent imaging was done using an IVIS Spectrum with excitation of 480 and emission of 570. The region of interest (ROI) of each plate was mapped and the fluorescence quantified using the instrument software (**Fig. 1**). In each well, multiple concentric circles were placed at 3, 6, and 10 mm out from the injection site.

**Figure 1:**
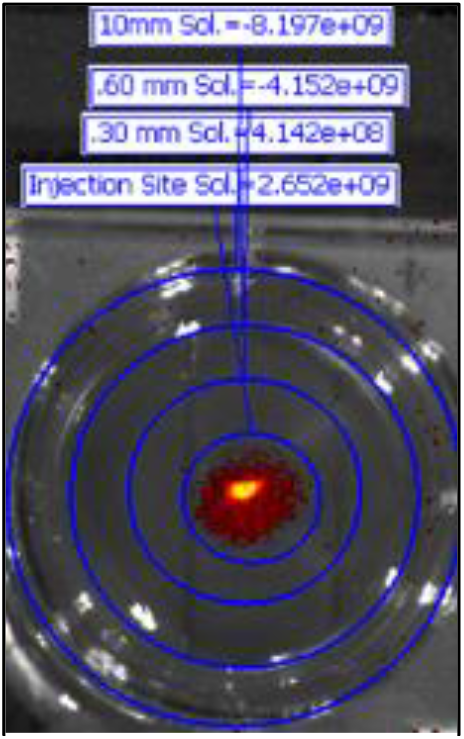
Example of ROIs used for quantifying distribution.

For one group a 1 mg/mL solution of DOX in DMSO and 1 mg/mL DOX in a 10% amygel was produced to be used to measure the distribution. Experiments were performed in triplicate and the data was averaged. The top row in the 6 well plate was each injected with 10µL of a 1 mg/mL DOX in DMSO solution. The bottom row was each injected with 10µL of a 1 mg/mL DOX in 10% amylopectin gel.

## Results

### Agarose Brain Phantom Allows for Fluorescent Observation of DOX *In Vitro*

The utilization of 0.6% agarose gel has been previously established as structurally similar to healthy brain tissues [18]. Placing degassed agarose gel into a polystyrene plate resulted in a completely clear system (**Fig.2**, top left image). When the agarose gel filled plate was imaged on the IVIS with the settings ideal for DOX, there was no background noise detected within the gel and minimal emission on the exterior edges of the plate (**Fig.2**, top left image). DOX was injected into the center of each well (**Fig.2**, top middle image), and was immediately visualized on the IVIS. This time zero image showed that the DOX was easily identifiable within each well (**Fig.2**, top right image).

**Figure 2:**
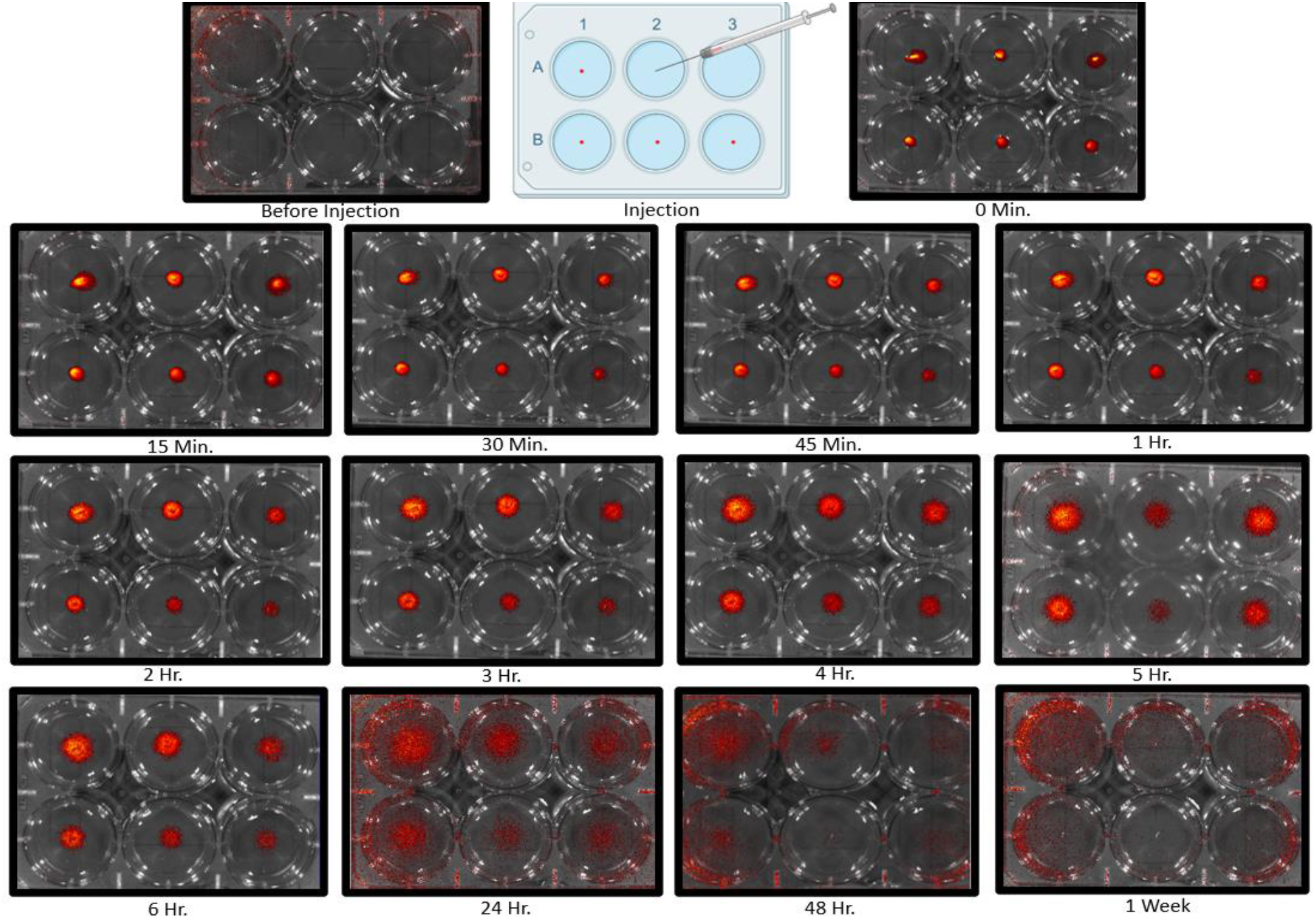
Fluorescent Imaging of DOX Diffusion Over One Week. Top left image is an empty agarose phantom with the top right image being the phantom immediatley post-injection. The top 3 wells in each plate was injected with DOX loaded into the amylose gel and the bottom 3 wells were injected with DOX in solution.

### Quantification of Emissions Demonstrate Gradual Diffusion of DOX from Solution and Gel

At 0 minutes post injected, for DOX in solution there was 3.46×10^9^, 9.17×10^9^, 1.33×10^9^, 2.95×10^9^ [p/s]/[µW/cm^3^] detected in the injection site, 3mm, 5mm, and 10mm ROIs respectively (**Fig. 3A**). At the same time, for DOX in the amylose gel there was 2.53×10^9^, 7.92×10^9^, 1.21×10^9^, 2.81×10^9^ [p/s]/[µW/cm^3^] detected in the injection site, 3mm, 5mm, and 10mm ROIs respectively (**Fig. 3B**). At 5 hours post injection the 3mm ROI had a larger emission than the initial injection site, and point reached by the amygel at 6 hours. Over the first 24 hours, the DOX was detected at levels of 3.56×10^9^, 5.91×10^9^, 6.79×10^9^, 9.03×10^9^ [p/s]/[µW/cm^3^] and 2.80×10^9^, 4.44×10^9^, 5.05×10^9^, 6.99×10^9^ [p/s]/[µW/cm^3^] for solution and gel. By 24 hours, both groups showed a notabley higher level of DOX in the 10mm ROI. In the 24, 48, and 1 week time points, there was an increase in background noise detected in the peripheral, focused within the plastic of the plate.

**Figure 3.**
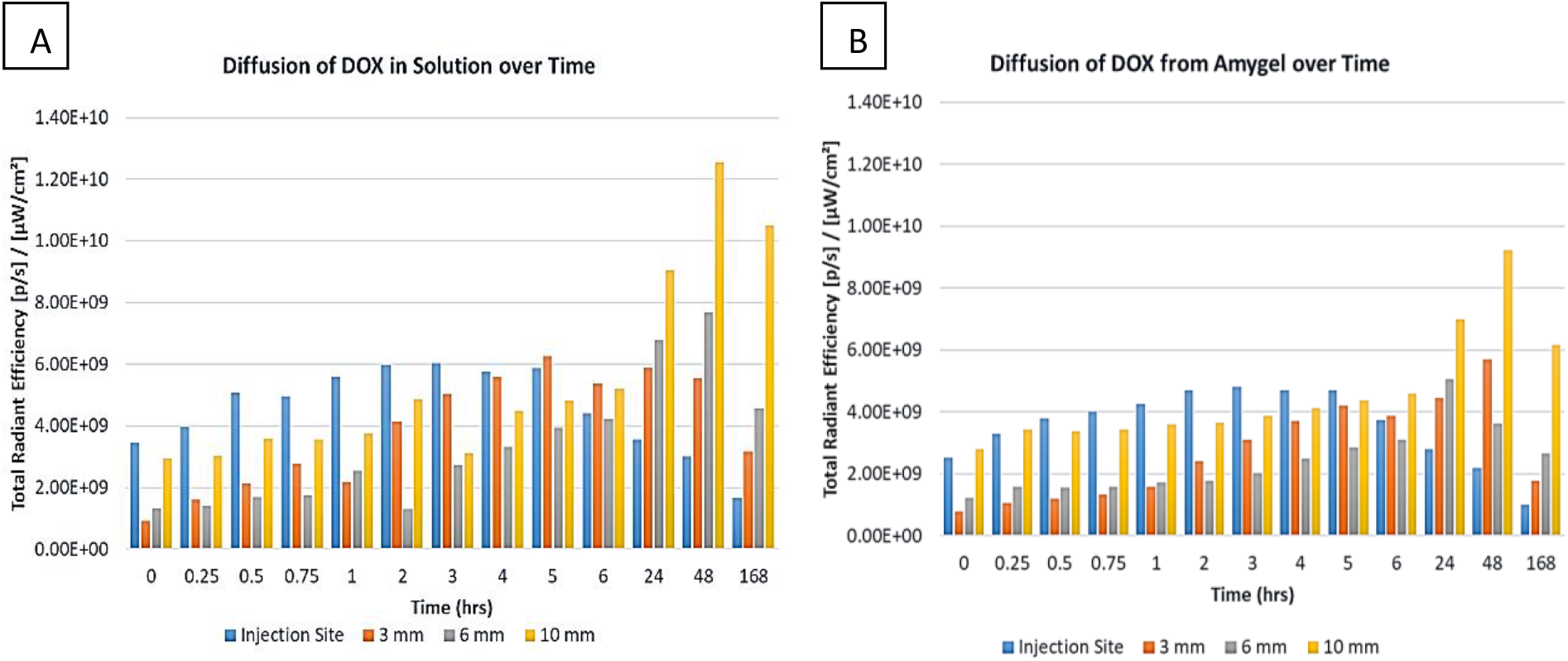
Intensity of DOX across diffused regions. Diffusion of DOX dissolved in solution (A) and DOX loaded amylose gel (B) quantified as radiant energy emitted in the concentric circle ROIs.

## Discussion and Conclusion

This study demonstrates the feasability of using an agarose gel tissue phatom as a mimic for healthy brain tissue to model the diffusion of a drug from the injection site. Over the past century, we have made strides in developinng chemotheraputic drugs. However, there remains a gap in having the pre-clinical cytotoxic sucesses translate to patient survival [3, 19]. Through this pilot experiment compairing the distribution of DOX susposended in solution and DOX loaded into a a 10% starch hydrogel we have shown that this testing platform allows for the monitoring of diffusion. The DOX appeared to be stable in the system over the first 2 days, and with general decreases in emission, it is a possibility that the drug began to degrade during days 2-7 (**Figs. 2-3**). DOX released from 10% Amygel diffused at a slower rate than the DOX solution (**Fig.3**). Compared to other chemotherapy agents, which are only able to penetrate microns into tissue [19-21], the DOX in both solution and Amygel was able to diffuse greater than 10 mm from the hydrogel implant.

To optomize this system, an addition of drug extract the agarose phantom for quantification of the amount of drug present using HPLC would increase the precision of diffusion and drug stability interpretation. Pairing this platform with *in vivo* data of drug penetration through murine tumor models would further the understanding of drug diffusion. These results demonstrate that this platform allows for the analysis of drug diffusion through brain tissue. Overall, these results illustrate a novel platform that can be paired with *in vivo* pre-clinical studies to optomize dosing of local drug delivery methods.

